# “Antimicrobial Properties and Physical Characteristics of Essential Oil Extracted from *Artemisia herba alba* Collected in El Bayadh, Algeria”

**DOI:** 10.1101/2023.04.13.536798

**Authors:** Mustapha mahmoud Dif, Djaadane Fatima zohra

## Abstract

*Artemisia herba-alba* is a white mugwort harvested in the Bougtobe region of the El-Bayad province. Essential oils were extracted from the plant using hydrodistillation. The antibacterial study of *Artemisia herba-alba* essential oils revealed a significant inhibitory action on the growth of tested microorganisms including *Bacillus cereus, Staphylococcus aureus*. These results demonstrated significant activity compared to the positive control penicillin, amoxicillin, and spiramycin. Overall, the results showed that the tested essential oil possesses interesting antimicrobial activities, and therefore, opens up new perspectives in the field of natural applications as a promising alternative to chemical products and for industrial exploitation.

## Introduction

*Artemisia herba alba* is a plant rich in secondary metabolites that offer medicinal properties. These metabolites include volatile constituents such as essential oils, and non-volatile constituents such as flavonoids and sesquiterpene lactones. The oil is qualitatively and quantitatively diverse, and several scientific studies have demonstrated its effectiveness as an antidiabetic, antihumanicide, antiparasitic, antibacterial, antiviral, antioxidant, antimalarial, antipyretic, antispasmodic, and antihemorrhagic agent (BOUDJELAL, 2013; Mohamed et al., 2010).

The antimicrobial properties of aromatic and medicinal plants have been known since ancient times. However, it was not until the early 20th century that scientists began to take an interest in them. Essential oils were found to inhibit and destroy all present microorganisms, ensuring virtually infinite preservation of the body. In old medical books, aromatic resins or essential oils were the active ingredients found in various plant drugs with significant antiseptic properties. In more recent works, the use of essential oils in aromatherapy offers a perspective of an alternative to synthetic drugs (Yano et al., 2006)

The objective of our work is to study the physicochemical characteristics of essential oils from *Artemisia herba alba* leaves and evaluate their antimicrobial activity.

## Materials and methods

### Study area

The Bougtob region covers an area of 2017.60 km² and is located in the northwestern part of the El-Bayadh province. The vegetation that grows in these arid zones has been used since ancient times as a source of food for wildlife and domestic animals, as well as for other purposes such as medicinal plants (Regagba, 1999).

### Sampling method

After harvesting, *Artemisia herba-alba* plant material (Figure1) is cleaned (free from debris), spread on paper, and allowed to dry at room temperature in a ventilated room, protected from humidity and light. This step is continued until the weight is stabilized. Once dried, the sample is ground and stored in glass containers tightly sealed and protected from humidity until the time of extraction.

**Figure1:**
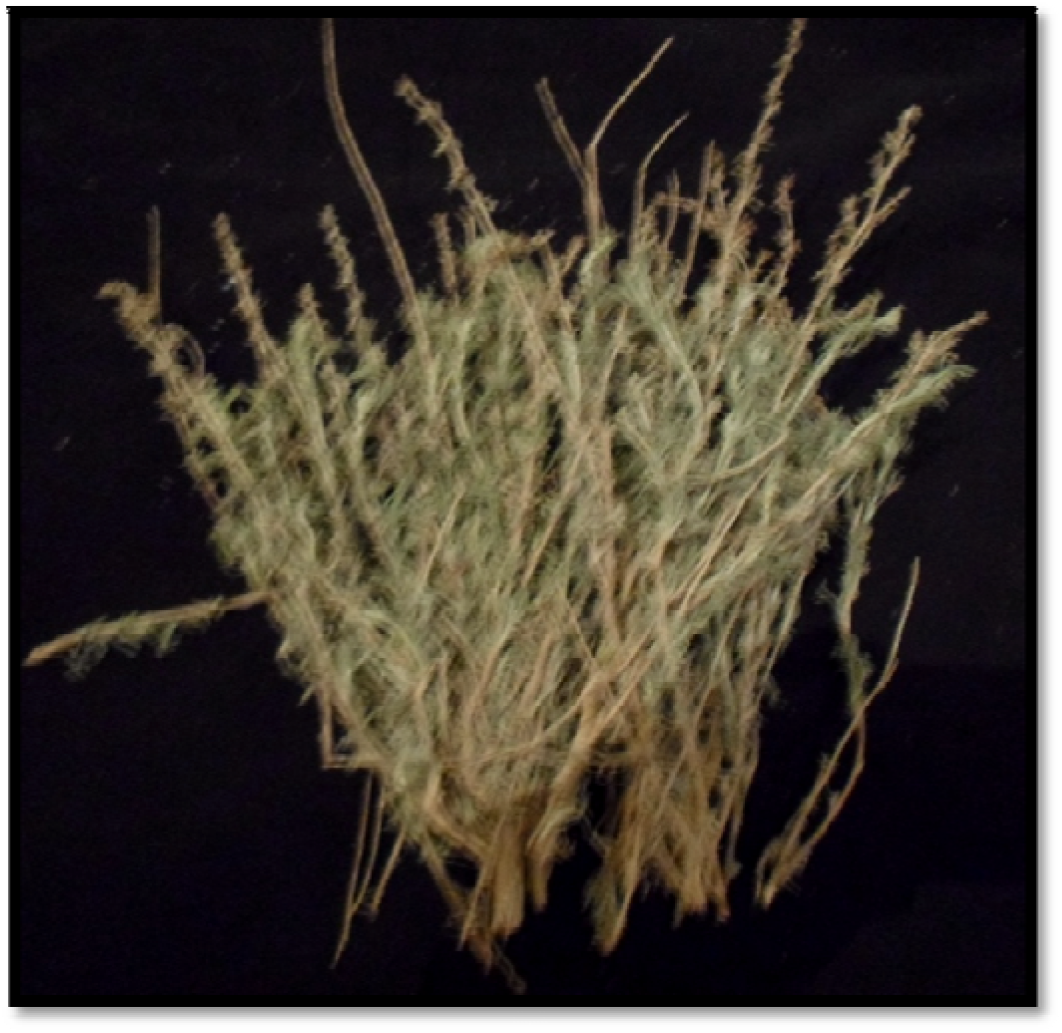
Presentation of *Artemisia herba alba* sample from Bougtob region

### The extraction of essential oil

The essential oil of *Artemisia herba-alba* was obtained by hydrodistillation, using a hydrodistillator. The setup consists of a 1-liter glass flask, which can hold 30g of plant material (aerial part of the plant) added with 500ml of water, placed above a heating mantle. The glass flask is topped with a column that communicates with a condenser, allowing the condensation of water vapors loaded with droplets of essential oil, which are then collected as distillate in a glass separatory funnel

### Physicochemical Characteristics of Essential Oils

Essential oil yield is expressed as a percentage and calculated by dividing the weight of extracted oil by the weight of plant material used. To assess the purity of an essential oil, physical characteristics such as the density and refractive index can be measured. Moisture content can be determined by calculating the percentage of water content before and after drying the sample, while pH can be measured using pH paper. The density of essential oil is determined by dividing the mass of a volume of oil by the mass of an equivalent volume of water at the same temperature. Refractive index is determined using a refractometer, by first passing a stream of water through the instrument to maintain it at the measurement temperature and adjusting it to give a refractive index of 1.3330 for distilled water at 20°C. The sample is then heated to the measurement temperature and the angle of incidence and refraction of a light ray passing from air into the essential oil are measured.

### Antimicrobial activity

#### Microbial material

Three bacteria: *Bacillus cereus, Staphylococcus aureus*, and *Staphylococcus hominis*

Two bacterial strains from the American Type Culture Collection (ATCC) were used in this study: *Bacillus cereus* ATCC and *Staphylococcus aureus* ATCC 25923.

Staphylococcus hominis from Spain, reference CECT 240 (Algerian CRAPC reference 18ESL)

### Preparation of microbial suspensions (inoculum)

The tested microbial strains are sub-cultured in test tubes containing approximately 5 mL of nutrient broth (NB) and incubated for 24 hours at 37°C to obtain a young culture. After homogenizing the microbial suspensions, their opacity should be equivalent to 0.5 McFarland or an optical density of 0.08 to 0.1 read at 620 nm. This density is adjusted by adding culture medium (NB) if it is too high. Then, the suspensions are streaked on petri dishes containing nutrient agar to verify their purity at 37°C for 24 hours

### Preparation of essential oil concentration

To prepare the decimal dilutions, Tween 20 was used to solubilize the essential oil, which has no antimicrobial activity.

Essential oils are generally very poorly soluble in water, which makes their biological and pharmacological study difficult. To solve this problem, several authors suggest the use of organic solvents such as acetone, DMSO or the use of emulsifying agents (surfactants such as Tween 20 or Tween 80) to help the essential oil solubilize in the culture medium (Duraffourd and Lapraz, 2002).

### Solid medium techniques: Vincent’s method

The antibioaromatogram technique or disk diffusion method (Dayal and Purohit, 1971) was used to evaluate the antimicrobial activity of essential oils. This test is performed by placing a sterile 6 mm diameter disk impregnated with a quantity of essential oil onto a pre-seeded solid medium with a microbial culture. After incubation, the results are read by measuring the diameters of the inhibition zones in millimetres (Chao et al., 2000; Andrews, 2001).

## Results and discussion

### Moisture content of the plant material

The determination of the moisture content *Artimisia herba alba* species from Bougtoub region is presented in Figure 2. The results showed a moisture content of 66.97%. This means that 33.02% represents the dry matter content that was actually used for the extraction of essential oils from this plant.

**Figure 2:**
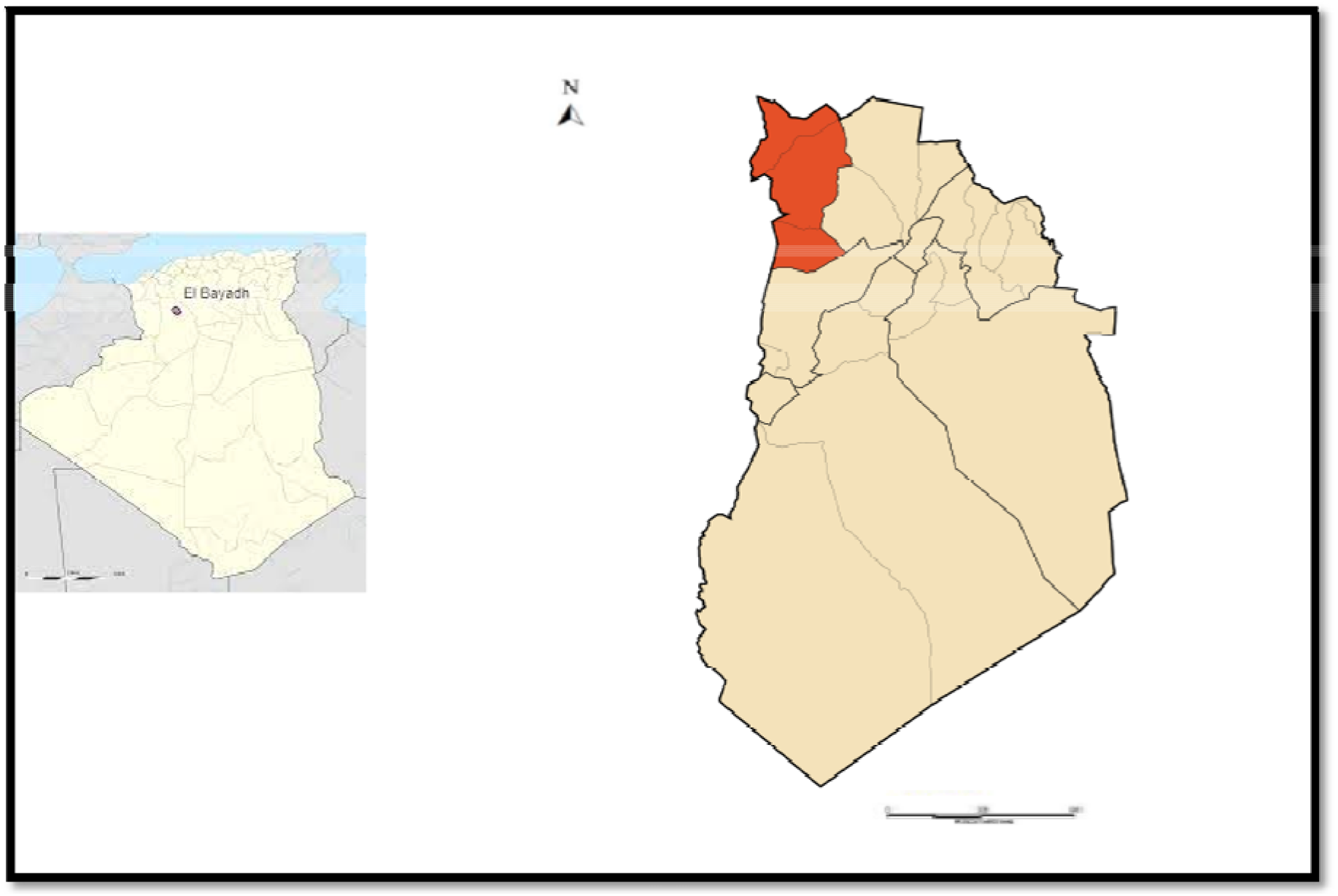
Presentation of the Bougtob region in relation to the El Bayadh province and Algeria

This moisture content is higher compared to what was reported in previous research on the Artimisia herba alba species from the Djelfa region, which was 48.53% (reference) (GOUDJIL Mohamed Bilal, 2016).

### Organoleptic quality

According to Afnor (2000), essential oils are usually liquid at room temperature and volatile, they are more or less colored and their density is generally lower than that of water.

The essential oil of *Artimisia herba alba* is a yellowish-green liquid with a strong and pleasant odour of white mugwort. These organoleptic properties are related to the climatic and soil conditions, the study region, the plant’s state, species, and the harvest period.

These organoleptic properties, such as colour, odour, and viscosity, can also be relevant for judging the quality of oil.

### Extraction yield

The extraction yield of *Artimisia herba alba* (Table1) species from the Bougtoub region was R% = 1.02%. It depends on several factors such as the species, the region of the harvest, the harvest period, the climatic and edaphic conditions of the study region, the plant’s condition, cultural practices, and extraction technique, among others (reference). The main conditions required for profitable essential oil production are good plant material, plant variety, soil, distillation equipment, and climate (Smallfield, 2001).

**Table 1:** the physicochemical characteristics of Arte essential oil

|  |  |
| --- | --- |
| Humidity level | 66.97 % |
| The yield | 1.02%. |
| The refractive index (T =20°C) | 1,471 |
| the relative density (T =20°C) | 1.12 |

For the *Artemisia* species, our yield is average compared to 18 Tunisian white wormwood provenances (0.68% - 1.93%) (Zouari, et al, 2009). The same intraspecific variation of white wormwood yield was noted in Spain (0.41% - 2.30%) for 16 samples from 4 provenances (Salido, 2004). The highest value was obtained with the *Artemisia herba alba* species with a yield of 1.7%.

### Refractive Index

The refractive index measured for our samples(Table1) is 1.471 for *Artimisia herba alba* oil at temperature T = 20°C. Therefore, a high refractive index indicates the presence of double bonds. The value is higher than the refractive index of water at 20°C (1.333) (Afssaps, 2008).

Our result is higher than that reported by Benjilali and Richard (1980), who showed that the refractive index of 24 white wormwood essential oil samples from different regions of Morocco varies from 1.4562 to 1.4696. This could be explained by the nature of the essential oil composition. This index varies according to the chemical composition, density, and increases with unsaturation or the presence of secondary functions (Boukhatem et al., 2010).

### Density

We observed that the relative density of the essential oil of *Artemisia herba alba* (Table1) is 1.12 at a temperature of 20°C. This physical characteristic is generally used in the classification of essential oils and is a crucial step but not sufficient to characterize them fully. It is, therefore, necessary to supplement it with other analyses. This parameter is related to the chemical composition of this oil, which is affected by a large number of factors such as the phenotype, harvesting time, type of soil, storage, extraction process, and conditions (Heidari, et al., 2008).

### Antimicrobial activity

The purpose of an antibiogram is to determine the Minimum Inhibitory Concentration (MIC) of a bacterial strain against various antibiotics.

The essential oil of *Artemisia herba-alba* plant reacted positively to the tested microbial strains. Large variations in inhibition zone diameters were observed, ranging from 7 to 28 mm. The most effective synthetic antibiotic (Table2) was found to be spéramicin, with inhibition zone diameters of 22, 21, and 25 mm for *Bacillus cereus, Staphylococcus aureus*, and *Staphylococcus hominis*, respectively. Amoxicillin also showed significant efficacy with inhibition zone diameters of 7, 22, and 16 mm, respectively. Penicillin, on the other hand, resulted in inhibition zone diameters of 10, 15, and 12 mm, respectively, classifying Staphylococcus in the group of sensitive microorganisms.

**Table 2:** Inhibitory effect of the synthetic antibiotic on the cultures: *Bacillus cereus, Staphylococci aureus and Staphylococuss hominis* (expressed in diameter of inhibition mm)

|  | penicillin | amoxicillin | spermamicin |
| --- | --- | --- | --- |
| C1: *Bacillus cereus* | 10 | 7 | 22 |
| C4: *Staphylococci aureus* | 15 | 22 | 21 |
| C7: *staphylococci hominis* | 12 | 16 | 25 |

Our results show a high variability in the bacteriostatic qualities of the oil against different strains. The essential oil’s activity varies from one strain to another, which can be observed in the inhibition zone diameters, indicating that certain strains are more sensitive than others (Ponce et al., 2003; Rota et al., 2008).

The antibacterial study of Artemisia herba-alba essential oils revealed a strong inhibitory effect on the growth of the tested microorganisms *Bacillus cereus, Staphylococcus aureus*, and *Staphylococcus hominis* showed in table 3

**Table 3:** Inhibitory effect of artemisia herba alba essential oil on crops: *Bacillus cereus, Staphylococci aureus and Staphylococuss hominis* (expressed in diameter of inhibition mm)

|  | Essential<br>oil mother<br>dilution | Dilution<br>10<sup>-1</sup> | Dilution<br>10<sup>-2</sup> | Dilution<br>10<sup>-3</sup> | Dilution<br>10<sup>-4</sup> | Dilution<br>10<sup>-5</sup> |
| --- | --- | --- | --- | --- | --- | --- |
| *Bacillus cereus* | 8.25±0.5 | 8.25±0.5 | 8.5±0.58 | 7.33±0.58 | 9.25±0.96 | 7±0 |
| *Staphylococci aureuse* | 8.25±2.8 | 8.25±0.95 | 8.5±0.57 | 10±0.81 | 7±0 | 7±0 |
| *Staphylococci hominis* | 8.25±0.5 | 9.25±0.5 | 7.25±0.5 | 9.75±0.5 | 8.25±0.5 | 8.25±0.5 |

These differences in microorganisms’ sensitivity to *Artemisia herba-alba* essential oil may be explained by the quantity and quality of bioactive molecules, the nature and composition of the cell wall, and the cell’s enzymatic system’s strength, which controls its metabolism. The antimicrobial activity of the essential oil of *Artemisia herba-alba* against S. aureus can be attributed to the combination of different components present in this oil (Bertella et al., 2018).

## Conclusion

The aim of our study was to investigate the antimicrobial properties of *Artemisia herba alba* essential oils and determine the physical quality of its essential oil extracted by hydrodistillation, as well as to highlight some of its biological properties. The obtained essential oil is yellow-green in colour with a strong odour and a liquid appearance. The determination of crude extract yields showed a significant yield of 1.02%, which is an essential characteristic when controlling, commercializing, or demonstrating potential specificity. The in vitro study of the inhibitory power of oils demonstrated a powerful and interesting effect on the majority of the tested strains, confirming the important and well-known role of these secondary metabolites in protecting against antibacterial effects of essential oils, as tested by the disk diffusion method. Moreover, the study of the biological activity on different pathogenic strains demonstrated good antimicrobial activity of our oils in vitro against tested bacterial strains such as B. cereus, S. aureus, and S. hominis. The antibacterial effect is proportional to the concentration of the essential oil. This study can be considered an important source of information on the antibacterial properties of *Artemisia herba alba* essential oils, which are a potential source of diverse compounds with biological activities, justifying their use in food and traditional medicine for the treatment of various pathologies and the development of new natural antibacterial agents.

